# Beneficial bacteria activate type-I interferon production via the cytosolic sensors STING and MAVS

**DOI:** 10.1101/792523

**Authors:** Jorge Gutierrez-Merino, Beatriz Isla, Theo Combes, Fernando Martinez-Estrada, Carlos Maluquer de Motes

## Abstract

Type-I interferon (IFN-I) cytokines are produced by innate immune cells in response to microbial infections, cancer and autoimmune diseases. These cytokines trigger protective responses in neighbouring cells through the activation of IFN-I stimulated genes. One of the most predominant pathways associated with IFN-I production is mediated by the cytosolic sensors STING and MAVS, intracellular adaptors that become activated in the presence of microbial nucleic acids in the cytoplasm, leading to IFN-I production via TANK-binding kinase (TBK)-1 and IFN regulatory factors. However, the role of these sensors in responses induced by beneficial microbes has been relatively unexplored. Here we have screened 12 representative strains of lactic acid bacteria (LAB), a group of beneficial microbes found in fermented food and probiotic formulations worldwide, for their ability to trigger IFN-I responses. Two isolates (*Lactobacillus plantarum* and *Pediococcus pentosaceus*) induced an IFN-I production that was significantly higher that the rest, both in macrophage cell lines and human primary macrophages. This response correlated with stronger interaction with macrophages and was susceptible to phagocytosis inhibitors, suggesting bacterial internalisation. Accordingly, macrophages deficient for STING and, to a lesser extent, MAVS failed to respond to the two LAB, showing reduced TBK-1 phosphorylation and IFN-I activation. Furthermore, LAB-induced IFN-I was biologically active and resulted in expression of interferon stimulated genes, which was also STING- and MAVS-dependent. Our findings demonstrate a major role for STING in the production of IFN-I by beneficial bacteria and the existence of bacteria-specific immune signatures, which can be exploited to modulate protective responses in the host.

## Introduction

Humans live in symbiosis with beneficial microbes that inhabit different parts of the body, including the gastrointestinal and respiratory tracts (1, 2). Beneficial microbes are composed of commensal bacteria that are critical to maintain homeostasis due to their involvement in appropriate host immune responses. One of the most significant bacterial groups associated with host beneficial properties are lactic acid bacteria (LAB) (3), typical commensals that play an important role in activating epithelial cells, antigen-presenting cells and phagocytes (4). Innate immune cells recognize LAB via the ligation of microbe-associated molecular patterns (MAMPs) to pattern recognition receptors (PRRs) such as Toll-like receptors (TLRs) and Nucleotide-binding oligomerization domain (NOD)-like intracellular receptors (5). Some MAMPs of LAB include pili (fimbriae), peptidoglycans, lipo-teichoic acids, exopolysaccharides and many other components of the cell wall and membrane (4). Upon MAMP recognition PRRs activate an intracellular signalling cascade that converges on the pro-inflammatory transcription factor nuclear factor kappa B (NF-κB), which is crucial in regulating anti-microbial responses in the mucosa (6). A few studies have also reported that LAB are capable of inducing protective pro-inflammatory responses that depend not only on NF-κB activation but also on interferon (IFN) regulatory factors (IRFs) (7, 8). The specific mechanisms that LAB utilize to activate IRFs are unknown but it has been reported that the production of type-I interferons (IFNs) requires the presence of TLRs in endosomal compartments, including TLR2 and TLR3 (7, 8).

Type-I IFN (IFN-I) such as IFN-α and IFN-β are polypeptides that innate immune cells secrete in response to infectious agents, particulary viral pathogens (9). IFNs are important to induce antimicrobial responses in an autocrine and paracrine manner via the synthesis of interferon stimulated genes (ISGs), and to modulate innate immune responses that result in a balanced natural killer cell response and effective antigen presentation to activate immunological memory through specific T and B cells responses. IFN-I can also display protective roles against bacterial infections, cancer and autoimmune diseases (10). The intracellular adaptors MyD88 and TRIF are very well-known to activate IFN-I production via TLRs in response to infections (11). More recently, the cytosolic molecules stimulator of interferon genes (STING) and mitochondrial antiviral signaling (MAVS) have been identified as potent IFN-I inducers in response to nucleic acids (12). STING is an endoplasmic reticulum-localized transmembrane protein that becomes activated in the presence of cytosolic DNA and the production of cyclic 2’,3’ GMP-AMP (cGAMP) by the cGAMP synthase (cGAS) (13-16) In addition, STING can also recognise other cyclic dinucleotides (CDN) of prokaryotic origin (17). MAVS is a mitochondria-localized transmembrane protein that is activated by retinoic acid inducigle gene-I (RIG-I) and MDA-5, two cytosolic sensors of 5’ tri- and di-phosphate RNA and stretches of dsRNA (18-20). Both STING and MAVS coordinate the activation of TANK-binding kinase (TBK)-1 and IRF3, which translocates into the nucleus to initiate IFN production (21, 22).

STING and MAVS are essential to induce protective immune responses in the presence of cytosolic DNA and RNA such as those from viral infections or intracellular pathogenic bacteria (23, 24). However, the role of these cytosolic sensors in the interaction with beneficial bacteria has remained relatively unexplored. Heretofore, we know that STING and MAVS detect DNA and RNA of gut commensal bacteria in order to induce appropriate immune responses (25, 26). Very recently, Moretti *et al*. have also reported that CDNs produced by live gram-positive bacteria, either pathogenic or commensal, activate STING (27); and Spiljar *et al*. have shown that IRF3 activation might be critical in maintaining gut homeostasis (28).

Here we have investigated the abilities of LAB to induce activation of the inflammatory transcription factors NF-kB and IRF-3 in human macrophages and found that, although most strains induced NF-κB activation, some were able to trigger a significant production of IFN-I that was dependent on the bacterial viability and STING. Internalisation of the IFN-I-inducing LAB by human phagocytes resulted in TBK-1 phosphorylation and the subsequent expression of IFN-β and ISGs. Taken together, these findings show that STING has a major role in activating immune responses mediated by beneficial bacteria and that these immune responses are specific to each bacterial group and species. Such bacterial immune signatures might be of particular importance to trigger and modulate protective responses including those mediated by IFN-I in healthy and diseased individuals.

## Materials and Methods

### Ethics statement

All procedures with human blood samples have been approved by the Surrey University Ethics Committee in accordance with the Institutional Policy on the Donation and Use of Human Specimens in Teaching and Research and the national guidelines under which the institution operates. The blood protocol approval was under IRAS number 236477 and the sample storage was carried out under HTA licence 12365.

### Human peripheral blood mononuclear cells (PBMC) and macrophages

Whole fresh blood was collected in heparinized tubes from healthy individuals to carry out the phagocytosis assay. To generate macrophages monocytes were first isolated from buffy coats of the healthy blood donors by Lymphoprep (Axis-Shield) density gradient centrifugation followed by plastic adherence in RPMI containing 15% FCS and 1% Pen/Strep. The collected peripheral blood mononuclear cells (PBMCs) were incubated for 2h to remove nonadherent cells by washing with PBS. The adherent monocytes were then detached using 10 mM EDTA in PBS at room temperature and plated for macrophage maturation in RPMI with FCS and Pen/Strep for 7 days.

### THP-1 cell lines and macrophage differentiation

THP-1 cells were propagated in Roswell Park Memorial Institute (RPMI) 1640 (Life Technologies) supplemented with 15 % foetal bovine serum (FCS, Seralab) and 1 % Penicillin/Streptomycin (Pen/Strep, Life Technologies) at 37 °C in an atmosphere of 5 % CO2. THP-1 stocks were prepared in FCS containing DMSO at 10 % and stored at -80°C. THP-1-κB-GLuc (Lucia™ NF-κB) express and secrete Gaussia luciferase (GLuc) under the control of a synthetic 5x-κB promoter (Invivogen. THP-1-IFIT1-GLuc express and secrete Gluc under the control of the promoter of the IRF-3-dependent gene IFIT1 and were a gift from Veit Hornung (29). THP-1-IFIT1 deficient for STING or MAVS, and its corresponding control, were kindly provided by Greg Towers and have been previously described (30). THP-1 monocytes were differentiated into macrophages by seeding 5×10^4^ cells per well (or 10^6^ for immunoblotting) in 96-well plates containing RPMI supplemented with phorbol 12-myristate 13-acetate (PMA) at 20 ng/mL for 48 h as previoulsy described (31). After the 48 h incubation medium was replaced for RPMI containing 2 % FCS and 1 % Pen/Strep and macrophages exposed to LAB. PMA was from Santa Cruz Biotechnology and kept in DMSO at 10 mg/mL.

### Growth and maintenance of LAB

The LAB selected for this study are shown in Table 1 and are reference strains with accessible peer-reviewed publications and/or genome sequences with the exception of *Pedioccocus acidilactici* (isolate E1) and *Lactobacillus kunkeei* (isolate O29). The genomes of the newly-characterized isolates E1 and O29 and their corresponding sequencing reads, assemblies and metadata have been uploaded onto Genbank in BioProject PRJNA544274. All the isolates were grown in MRS broth (Oxoid) at 37°C without any aeration for 24 h, and maintained as -80°C frozen stocks in their appropriate media with the addition of 15% glycerol.

**TABLE 1.** Lactic Acid Bacteria (LAB) isolates used in this study.

| Species | Strain | Internal Code | Source | Reference | Donor |
| --- | --- | --- | --- | --- | --- |
| *Enterococcus faecalis* | NCTC 8213 | EF | Human babies' faeces | (57) | Surrey Culture Collection |
| *Lactobacillus casei* | DN-114001 | LC | Dairy product | Commercial | Isolated from Actimel, Danone |
| *Lactobacillus kunkeei* | O29 | LK | Bee pollen | This study |  |
| *Lactobacillus plantarum* | WCFS-1 | LP | Human saliva | (58) | Prof M Kleerebezem, Wageningen University, the Netherlands |
| *Lactobacillus sakei* | Lb790 | LS1 | Meat | (59) | Dr L Axelsson, Nofima AS, Norway |
| *Lactobacillus sakei* | L45 | LS2 | Norwegian fermented dried sausage | (60) | Prof I Ness, Norwegian University of Life Sciences |
| *Lactococcus lactis* | BBS4 | LL1 | Spanish fermented dried sausage | (61) | Prof PE Hernandez Cruza, Universidad Complutense de Madrid |
| *Lactococcus lactis* | IO-1 | LL2 | Kitchen sink water | (62) | Prof H Yoshikawa, Tokyo University of Agriculture |
| *Pediococcus acidilactici* | PAC1.0 | PA1 | Fermented food | (63) | Prof J Kok, Groningen University, the Netherlands |
| *Pediococcus acidilactici* | E1 | PA2 | Raw honey | This study |  |
| *Pediococcus pentosaceus* | NCTC 990 | PP | Dried beer yeast | (64) | Surrey Culture Collection |
| *Streptococcus thermophilus* | DN-001116 | ST | Yoghurt | Commercial | Isolated from a Danone yoghurt |

### Bacterial challenge

The macrophages derived from the pNF-κB-GLuc and pIFIT1-GLuc monocytes were exposed to inactivated and viable LAB cells. The inactivated cells were used as heat-treated (70 °C, 2 h) LAB pellets at a ratio of 100 cells per 1 macrophage and incubated for 24 h, while viable cells were incubated simultaneously with the macrophages for 2h in RPMI withouth Pen/Strep at differnt doses as indicated in the figure legends before being replaced for fresh RPMI with antibiotics to incubate for a further 22 h. To measure Gluc activity, media was collected from the macrophage cultures and transferred to white-bottom 96-well plates, which were read in a Clariostar plate reader (BMG Biotech) in the presence of 2 μg/mL of coelenterazine (NanoLight Technology). LPS at 0.2 μg/μl (Sigma Aldrich) was used as a control for the activation of NF-κB and IFIT1. Activation of NF-κB or IFIT1 was calculated as a fold increase ± SD over the measurements recorded for unchallenged macrophages.

### Bacterial intake in THP-1 macrophages

In order to estimate the number of LAB that are internalized by THP-1 macrophages we carried out an extracellular survival assay as previously described (32) with minor modifications. Macrophages were incubated with viable LAB at a ratio of 25 bacteria per phagocyte in RPMI containing 2 % FCS for 2h, at 37°C and 5% CO_2_. Prior to the LAB exposure macrophages were treated with the phagocytosis inhibitor cytochalasin D at a final concentration of 10 μM (Sigma-Aldrich). LAB were also incubated without macrophages under the same experimental conditions (with or without cytochalasin D). Supernatants were then collected at time points 0 and 2h to calculate and compare the number of bacteria that remain viable and detached from THP-1 macrophages. Bacterial enumeration was performed by quantitative plating of serial dilutions on MRS agar plates.

### Phagocytosis assay with human peripheral blood mononuclear cells (PBMC)

Approximately 10^6^ leukocytes were mixed 1:1 with 10 mM EDTA in PBS and challenged with LAB that were previously labelled with the FITC dye (Sigma), at a multiplicity of infection (MOI) of 25 bacteria per cell. The phagocytosis incubation was then carried out at 37°C in an orbital shaker for 1 h and subsequently treated with 1x RBC lysis solution (Biolegend) following incubation at room temperature for 15 min. The cells were washed twice with EDTA-PBS, resuspended in PBS and analyzed on FACS Celesta instrument (BD Biosciences). Flow cytometry analysis based on forward (FSC) and side (SSC) scatter was used to distinguish the main blood cell populations based on their size and granularity (lymphocytes vs. phagocytes), while the FITC channel was used to measure bacterial interaction with blood cells. The resulting SCC/FITC plots were then used to quantify the LAB intake by phagocytes.

### CD expression in phagocytes from human PBMCs

In order to monitor the expression of CD40 and CD64 in monocytes and neutrophils exposed to LAB we carried out the phagocytosis assay as described above but with some modifications. Leukocytes and bacteria were mixed at 1:100 ratios and incubated for 1 h. After phagocytosis incubation antibodies against human CD40-BV510 and CD64-PE/Dazzle (Biolegend), or the corresponding isotype controls, were added to the samples and incubated for 20 min at room temperature. The fixable viability dye Zombie NIR (Biolegend) was also used for a live/dead stain as per manufacturer’s instructions. Cells were then fixed with RBC lysis buffer and the CD expression compared to the matching isotype control and no-bacteria controls to calculate expression levels in each of the blood cells populations identified by FACS.

### Microscopy studies on monocyte-derived macrophages from human PBMCs

Monocyte-derived macrophages were challenged for 2 h with FITC-labelled LAB as described above for the THP-1 cells. After phagocytosis macrophages were fixed for 20min with 1% paraformaldehyde (PFA), washed with PBS without Calcium or Magnesium and stained with DAPI for imaging on an EVOS fluorescent microscopy.

### SDS-PAGE and immunoblotting

LAB-challenged THP-1 cells were lysed in radio-immuno-precipitation assay (RIPA) buffer supplemented with protease & phosphatase inhibitors (Roche) and benzonase at 250 U/ml (Sigma). Lysates were rotated for 30 min at 4 °C and subsequently denatured for 5 min at 95 °C in the presence of loading buffer and resolved by SDS-PAGE. The samples were then transferred to nitrocellulose membranes (GE Healthcare) using a Trans-Blot semidry transfer unit (Bio-Rad). Membranes were blocked in 0.1 % Tween/phosphate-buffered saline supplemented with 5 % skimmed milk (Sigma) and subjected to immunoblotting with the following primary antibodies at the indicated dilutions: phosphorylated TBK-1 Ser^172^ (Abcam; 1:5,000), TBK-1 (Abcam; 1:5,000) and α-tubulin (Upstate Biotech; 1:10,000). Primary antibodies were detected using IRDye-conjugated secondary antibodies in an Odyssey infrared imager (LI-COR Biosciences). Images were analyzed using Odyssey software.

### Quantitative RT-PCR

PMA-differentiated THP-1 cells were exposed to LAB for 8 h and cells were collected for RNA extraction using the High Pure RNA Isolation Kit (Roche Diagnostics Limited) following manufacturer’s instructions. RNA was reverse transcribed using the SuperScript ® III First-Strand Synthesis System from Invitrogen and the resulting cDNA diluted 1:5 in water. cDNA was then used as a template for real-time amplification on a QuantStudio 7 Flex Real-Time PCR system (Applied Biosystems) using SYBR Green Master Mix (Applied Biosystems) and specific primers for: TATA box Binding Protein (TBP) (forward, 5’ TGCACAGGAGCCAAGAGTGAA; reverse, 5’ CACATCACAGCTCCCCACCA), myxovirus resistance gene A (MxA) (forward, 5’ CCCCAGTAATGTGGACATCG; reverse, 5’ ACCTTGTCTTCAGTTCCTTTGT), human 2’5’-oligoadenylate synthetase 1 (OAS1) (forward, 5’ TGTGTGTGTCCAAGGTGGTA; reverse, 5’ TGATCCTGAAAAGTGGTGAGAG), and human IFNβ (described previously (33)) using three technical replicates of samples obtained from three biological replicates. Expression of each gene was normalized to the house-keeping gene TBP, and these values were then normalized to the non-stimulated control cells to yield a fold induction.

### ISRE-based IFN-I bioassay

Cell culture supernatants from LAB–challenged, PMA-differentiated THP-1 macrophages were collected after 24 h of incubation and transferred to 96-well plates containing HEK293T cells previously transfected for 24 h with 70 ng/well of a reporter plasmid expressing firefly luciferase (FLuc) under the control of IFN-stimulated responsive elements (ISRE) and 10 ng/well of a control plasmid expressing *Renilla* luciferase (RLuc) (Promega) using polyethylenimine (PEI; Sigma) at a ratio of 1:2 (μg of DNA:μl of PEI). After 10 h the cells were lysed in passive lysis buffer (Promega) and the FLuc and RLuc activities measured in the Clariostar plate reader (BMG Biotech). The FLuc/RLuc ratios were then calculated for each well and normalized to mock-infected THP-1 samples to be presented as a fold increase of ISRE activation.

### ELISA cytokine detection

Human TNF-α production was detected and quantified in the supernatants of LAB-challenged PMA-differentiated THP-1 cells using the eBioscience Human TNF-α ELISA Ready-SET-Go kit as indicated by the manufacturer’s instructions. The Human IFN-β Quantikine ELISA Kit (R&D systems) was used to detect and quantify IFN-β in supernatants from primary macrophages exposed to LAB for 2h. Supernatants were collected every 4 h for 12h after the 2 h phagocytosis

### Statistical analysis

Statistical analysis was performed using GraphPad Prism. Data are presented as means ± standard deviation (SD) and are representative of one experiment of at least three independent experiments. Data from experiments with human peripheral blood mononuclear cells are representative of two or three healthy donors and are mean with SD from two biological replicates. Statistical significance between one sample and its corresponding control was determined using the Student’s *t*-test and within a group of samples using one way ANOVA followed by Fisher’s Least Significant Difference (LSD) Test.

## Results

### LAB activate NF-κB or IFN-I responses in THP-1 macrophages

The ability of 12 LAB strains (Table 1) to trigger the activation of the inflammatory transcription factors NF-κB and IRF-3 was evaluated in human differentiated THP-1 cells. We employed 2 lines of THP-1 monocytes expressing GLuc under the control of either the NF-κB promoter or the promoter of the IRF-3-dependent gene IFIT1. The cells were differentiated for 48 h and subsequently exposed to live or dead LAB before GLuc activity was measured in the media and presented as a fold increase over non-stimulated macrophages (Fig. 1A). The majority of LAB species tested –*S. thermophilus, P. acidilactici, L. sakei, L. kunkeei, L. casei* and *E. faecalis*-induced significant NF-κB activation in THP-1 macrophages and, with the exception of *E. faecalis*, this activation was enhanced in response to live bacteria compared to inactivated bacterial cells. By contrast, we found that these isolates were poor inducers of IFIT1 activation either dead or alive. Interestingly, two bacterial species - *Pediococcus pentosaceus* (PP) and *Lactobacillus plantarum* (LP) - were able to induce a significant IFIT1 activation. This IFIT1 response was only observed with viable bacteria and was especially high with *L. plantarum*. Interestingly, neither PP nor LP enhanced NF-κB activation when used as viable bacteria. To visualize this contrasting behavior, we then plotted the recorded NF-κB vs IFIT1 activation for each LAB species and drew arbitrary cut-offs for NF-κB activation (∼20 fold) and IFIT1 (∼10 fold) based on the most common response observed in all bacterial species tested (Fig. 1B). As expected most LAB grouped in the top left quadrant (Q1) as powerful NF-κB agonists, but poor IFIT1 inducers. Strikingly, LP and PP appeared in the bottom right quadrant (Q4) as remarkably strong IFIT1 inducers, especially LP. The 2 *L. lactis* species occupied the bottom left quadrant (Q2) as intermediate NF-κB and IFIT1 inducers. We then performed dose-dependent exposures with LP and PP and showed that a challenge with 1 LP and 10 PP bacterial cells per macrophage was sufficient to trigger IFIT1 activation (Fig. 1C). The highest IFIT1 response was observed with a ratio of 25 bacteria per macrophage. Our screening therefore indicated that some LAB could activate responses converging on IFN-I production.

**Fig. 1.**
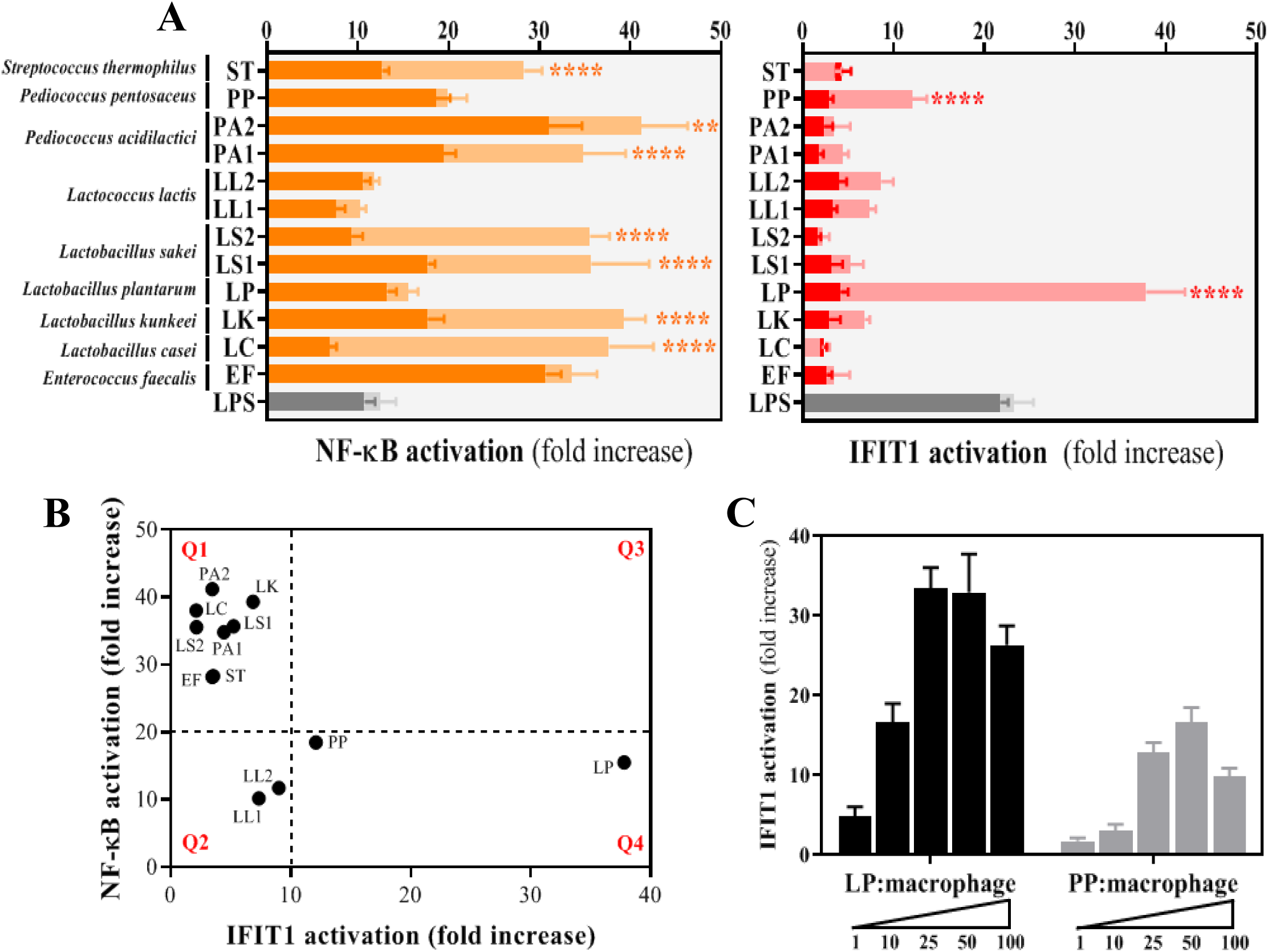
Out of the 12 LAB strains selected in this study, only viable cells of *Lactobacillus plantarum* (LP) and *Pediococcus pentosaceus* s(PP) sgnificantly uce IFIT1 activation in THP-1 macrophages in a dose-dependent manner. A. Response of THP-1 macrophages to LAB isolates as a measurement of NF-kB (orange) and IFIT1 (red) activation. The activation of NF-kB (or IFIT1) is presented as a fold increase over a non-stimulated condition using a PMA-differentiated pNF-kB-GLuc THP-1 reporter cell line (or pIFIT1-GLuc THP-1) that was exposed to LAB at a ratio of 1 macrophage per 100 inactivated bacterial cells (dark orange and red) or 25 viable bacteria (light orange and red). LPS was used as a positive control for the activation of NF-κB and IFIT1 under all experimental conditions (grey bars that are dark and light represent controls for experiments with inactivated bacterial cells and viable bacteria, respectively). For each of the isolates, a comparative statistical analysis was carried out between inactivated and viable cells using the Student t-test (**p < 0.01, ****p < 0.001). B. Correlation between NF-kB and IFIT1 activation in THP-1 macrophages exposed to LAB selected at a ratio of 1 macrophage per 25 viable bacteria. Activation is presented as a fold increase over a non-stimulated condition using PMA-differentiated THP-1 cells of pNF-kB-GLuc and pIFIT1-GLuc. Fold increases of 20 and 10 were selected as arbitrary thresholds for NF-kB and IFIT1 activation, respectively, to divide the illustration into 4 quadrants (Q1, Q2, Q3 and Q4). C. IFIT1 activation in THP-1 macrophages challenged with viable cells of LP (black bars) and PP (grey bars) at increasing macrophage:bacteria ratios of 1:1, 1:10, 1:25, 1:50 and 1:100. The activation is presented as a fold increase over a non-stimulated condition using a PMA-differentiated pIFIT1-GLuc THP-1 reporter cell line.

### Lactobacillus plantarum (LP) and Pediococcus pentosaceus (PP) activate IFN-I production

To address expression of IFN-I and verify the activation observed using luciferase reporters, we used a commercial ELISA against the NF-κB-dependent inflammatory cytokine TNF-α and an ISRE-based bioassay to quantify the presence of TNF-α and IFN-I in the supernatants of macrophages exposed to each of the LAB isolates (Fig. 2A and 2B, respectively). Apart from LP, PP and isolates of *L. lactis* (LL), the exposure of macrophages to all LAB isolates resulted in high amounts of TNF-α in the media, but very low levels of IFN-I. Conversely, LP and PP were capable of inducing a significant ISRE activation as indicative of effective production of IFN-I, especially in the case of LP (Fig. 2B). Similarly to the luciferase-based results, when the production of TNF-α and IFN-I were plotted we identified a strong IFN-I signature for LP that correlated with a very low impact on TNF-α (bottom right quadrant Q4 of Fig. 2C). PP showed a more moderate IFN-I signature and a slightly higher TNF-α production than LP (next to the bottom left quadrant Q2 of Fig 2C), but this was still much lower than that observed with most of the LAB isolates (top left quadrant Q1 of Fig. 2C). Taken together, our screens in human macrophages demonstrated that each LAB has the ability to stimulate innate immunity differently and that some are potent inducers of IFN-I responses.

**Fig. 2.**
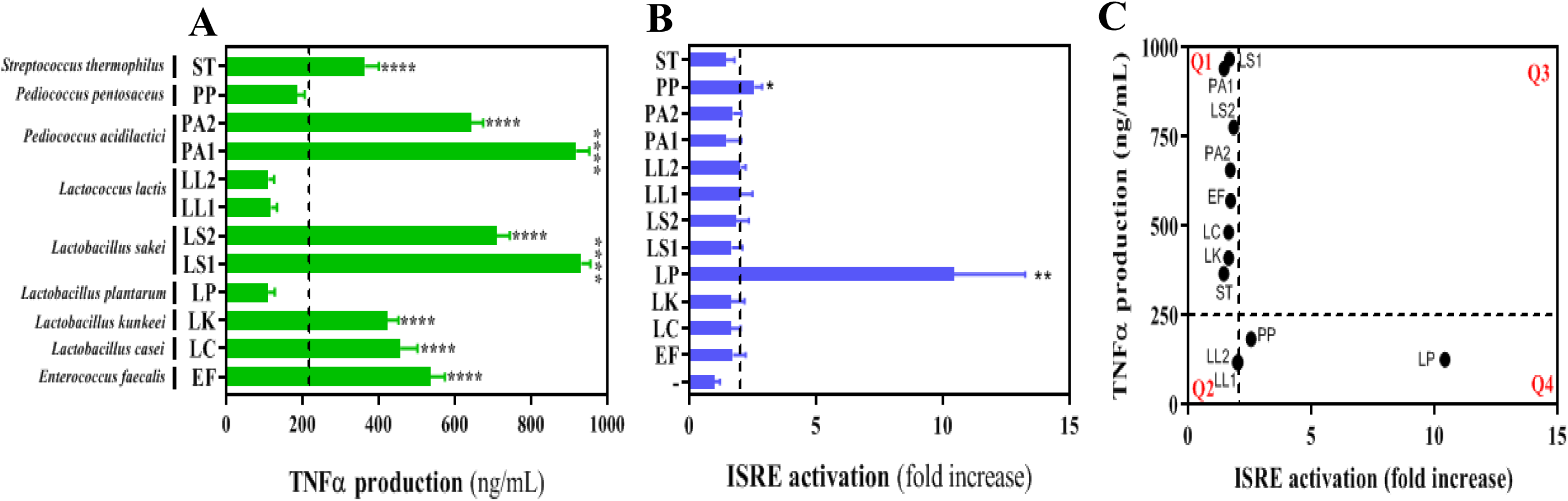
*Lactobacillus plantarum* (LP) and *Pediococcus pentosaceus* (PP) barely induce TNFa production in THP-1 macrophages but their influence on the E activation in these macrophages is significant. A. TNFa production in supernatants obtained from THP-1 macrophages exposed to LAB isolates at a ratio of 1:25. TNFa was detected and quantified as ng/mL using a commercial TNF-α ELISA. Isolates on the right hand side of the dotted line induce a significant production of TNFa by comparison with the isolates located on the left hand side. Differences between the isolates was carried out using one way ANOVA followed by Fisher’s Least Significant Difference (LSD) Test (****p < 0.001). B. ISRE activation from supernatants obtained from THP-1 macrophages exposed to LAB isolates at a ratio of 1:25. The ISRE activation is calculated as a fold increase using a pISRE-FLuc/RLuc reporter cell line after 10h of supernatant exposure. Isolates on the right hand side of the dotted line induce a significant ISRE activation by comparison with the isolates located on the left hand side Differences between samples including the cells stimulated with each of the isolates and the control (-, non-stimulated cells) was carried out using one way ANOVA followed by Fisher’s Least Significant Difference (LSD) Test (*p < 0.05, **p < 0.01). C. Correlation between TNFa production and ISRE activation in THP-1 macrophages exposed to each of the LAB selected in this study at a ratio of 1:25. A TNFa concentration of 250 ng/mL and an ISRE fold increase of 2 were selected as the arbitrary thresholds to divide the illustration into 4 quadrants (Q1, Q2, Q3 and Q4).

### Lactobacillus plantarum (LP) and Pediococcus pentosaceus (PP) activate responses associated with IFN-I production in human PBMCs

Next we sought confirmation of this LAB-induced IFN-I activation in human primary cells.We first employed an IFN-β ELISA to detect and quantify the presence of this cytokine in the supernatants of human monocyte-derived macrophages collected from PBMC and exposed to LP and PP for 2h. We observed a peak of IFN-β production 8 h post-challenge that decreased at 12 h (Fig. 3A). We then used flow cytometry to monitor the expression of the IFN-I-related markers CD64 and CD40 (34-36) in monocytes collected from PBMC from two healthy donors upon exposure to LAB. IFN-I expression has been shown to down-regulate CD64 (36) and upregulate CD40 in the presence of LAB (35). In agreement with this, we observed that only monocytes exposed to IFN-I-producing LP and PP showed a very significant reduction in CD64 expression (Fig. 3B) and an increase in CD40 expression (Fig. 3C). No significant changes were observed in the presence of *Lactobacillus casei* (LC), an isolate that showed a potent NF-κB response but poor IFN-I activation in our initial LAB screening. These findings therefore confirmed that LAB can trigger IFN-I responses in human primary immune cells.

**Fig. 3.**
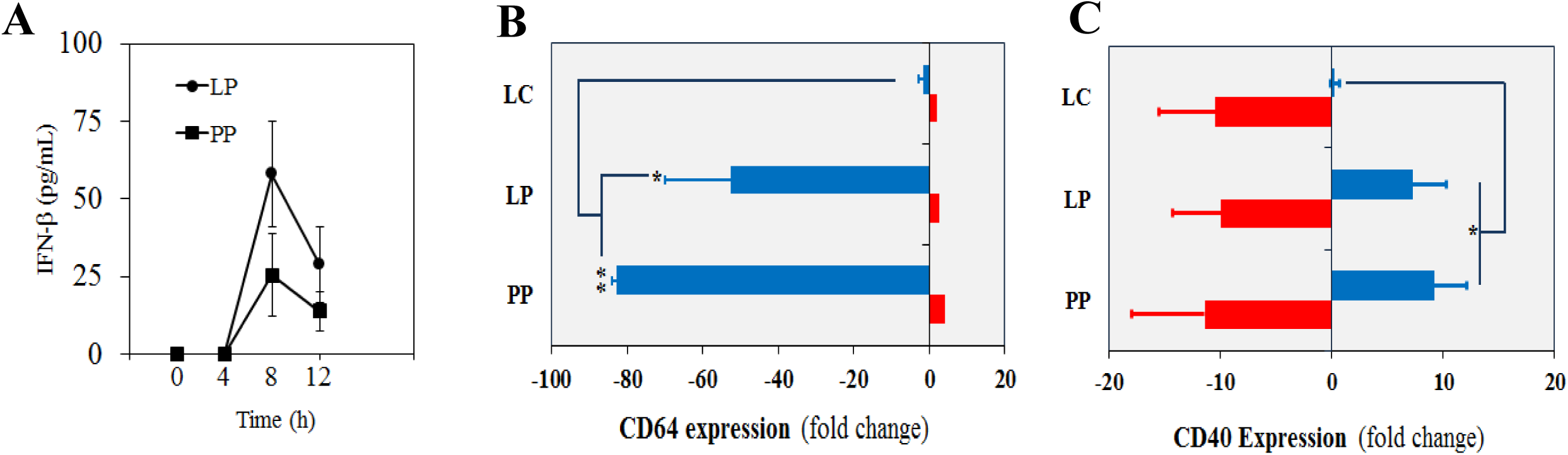
*Lactobacillus plantarum* (LP) and *Pediococcus pentosaceus* (PP) induce responses associated with IFN-I production in monocytes and macrophages m PBMCs. A. Production of IFN-b in supernatants obtained from primary macrophages after 0, 4, 8 and 12h of being exposed to LP and PP. The INF-b levels were calculated using the Human IFN-b Quantikine ELISA Kit. Data are representative of three healthy donors and are mean with SD from two biological replicates. B. Expression of CD64 in monocytes (blue) and neutrophils (red) exposed to LP and PP. The CD expression is represented as a percentage of fold change (increase or decrease) over a non-stimulated condition using unchallenged monocytes and neutrophils. *Lactobacillus casei* (LC) is included as a negative control (no IFN-I inducer). Data are mean with SD from two healthy donors and the comparative analysis was carried out with two way ANOVA and Tukey multiple comparison (* p<0.05, ** p<0.01). C. Expression of CD40 in monocytes (blue) and neutrophils (red) exposed to LP and PP. The CD expression is represented as a percentage of fold change (increase or decrease) over a non-stimulated condition using unchallenged monocytes and neutrophils. *Lactobacillus casei* (LC) is included as a negative control (no IFN-I inducer). Data are mean with SD from two healthy donors and the comparative analysis was carried out with two way ANOVA and Tukey multiple comparison (* p<0.05, ** p<0.01).

### LP and PP interact with or/and are internalized by human monocytes and macrophages

We next sought to determine the mechanism by which some LAB induce IFN-I production. First we examined whether LP and PP interact with human phagocytes –monocytes and macrophages-interact and/or phagocitose viable cells of LP and PP we carried out 3 independent phagocytic studies using PBMCs, macrophages differentiated from PBMCs and THP1 macrophages, at a ratio of 1 phagocyte per 25 bacteria. This ratio ensures a maximum activation of IFIT in phagocytes exposed to either LP or PP (Fig. 1C). Firstly, our phagocytosis assay with PBMCs showed that monocytes and neutrophils bind to or uptake LP and PP (Fig. 4A-B-C). By comparison with the NF-κB inducer *Lactobacillus casei* (LC), a significant number of monocytes and neutrophils phagocytose LP and PP (Fig. 4B-C). However, most of the phagocytes that interact with LP and PP are present in the populations to which these bacteria bind. Secondly, confocal microscopy allowed for the generation of images showing the detection of LP and PP inside PBMCs-derived macrophages (Fig. 4D). This demonstrated that LP and PP possess the capacity to enter human phagocytes. And thirdly, we observed that the number of LP and PP that remain inside (and/or attached to) THP1 macrophages is significantly higher in the presence of the phagocytosis inhibitor cytochalasin D following a 2h incubation in RPMI (Fig. 4E). We also recorded a decrease in bacterial viability after the 2h incubation, but with no significant differences in RPMI with or without cytochalasin D (Fig. 4F). Hence, these results strongly suggest that LP and PP interact and are internalized by human macrophages.

**Fig. 4.**
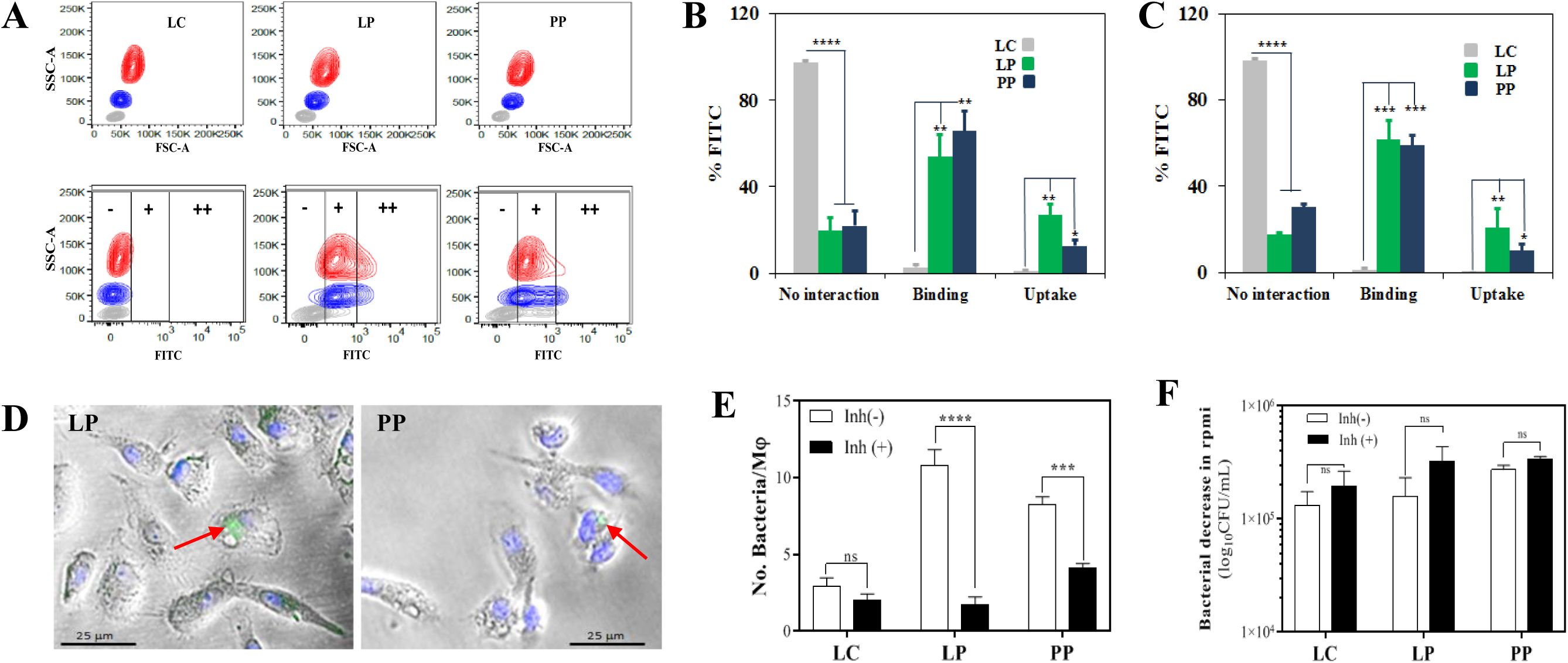
Human phagocytes interact with *Lactobacillus plantarum* (LP) and *Pediococcus pentosaceus* (PP) A. Uptake of LP and PP by monocytes and neutrophils from PBMCs of 2 healthy donors. Viable bacterial cells were labelled with FITC and incubated in whole human blood for 1h at a multiplicity of infection of 1:25. Blood cell populations were distinguished based on side scatter area (SSC-A) versus forward scatter area (FSC-A), and separated into lymphocytes (grey), monocytes (blue) and neutrophils (red). Phagocytic uptake was then observed in the FITC channel, where the intensity was divided into three subpopulations based on the positivity: no interaction (**-**), surface binding (**+**), and phagocytic uptake (**++**). *Lactobacillus casei* (LC) is included as a control of no phagocytic uptake. B. Percentage of monocytes in each of the three FITC subpopulations -no interaction, surface binding and phagocytic uptake-after exposure to LC (grey), LP (green) and PP (blue). The comparative analysis was carried out with two way ANOVA and Tukey multiple comparison *, p<0.05 **, p<0.01; ****, p<0.001). C. Percentage of neutrophils in each of the three FITC subpopulations -no interaction, surface binding and phagocytic uptake-after exposure to LC (grey), LP (green) and PP (blue). The comparative analysis was carried out with two way ANOVA and Tukey multiple comparison *, p<0.05 **, p<0.01; ***, p<0.005 ****, p<0.001). D. Confocal microscopy images showing the detection of LP and PP inside monocyte-derived macrophages (indicated with arrows). FITC and DAPI were used to label bacteria and the macrophage nucleus, respectively. E. Number of bacteria (LC, LP and PP) that remain inside (and/or attached to) THP1 macrophages after a 2h RPMI incubation in the absence (black) or presence (white) of the phagocytosis inhibitor cytochalasin D. The bacterial intake (and/or attachment) was estimated as the difference between number of bacteria per macrophage (Mj) that are present in the supernatants of THP1 macrophage cultures exposed to LP (or PP) at a ratio of 1:25 for 0 and 2h. Comparative analysis was carried out using the Student t-test (ns, no significant; ***p<0.005, ****p<0.001). F. Decrease in bacterial counts for LC, LP and PP after a 2h RPMI incubation in the absence (black) or presence (white) of the phagocytosis inhibitor cytochalasin D. The bacterial decrease was calculated as the difference between number of viable bacteria per mL after 0 and 2h of incubation and expressed as log_10_CFU/mL. Comparative analysis was carried out using the Student t-test (ns, no significant; ***p<0.005, ****p<0.001).

### STING and MAVS sense LP and PP to activate the IFN-I associated kinase TBK1

STING and MAVS are potent IFN-I inducers that respond to cytosolic PAMPs. Having observed that IFN-I-inducing LAB were internalised by macrophages, we explored whether these cytosolic sensors could account for the observed immune responses. We then used THP-1-IFIT1-GLuc cells deficient for STING (STING KO) or MAVS (MAVS KO) (30). We differentiated these cells into macrophages and challenged them with viable cells of PP and LP before measuring GLuc activity. As expected, control cells responded strongly to LP and IFIT1 activation reached 10 and 40 fold increase at 12 and 24 h after challenge. However, this response was completely abrogated in STING KO cells (Fig. 5A). The response in MAVS KO cells was also reduced, but it was less evident, being only statistically significant after 24 h of bacterial challenge. To verify the role of STING in LP-induced IFIT-1 activation, we examined the levels of phosphorylated TBK-1 in the lysates of cells exposed to different LP doses. Phosphorylated TBK-1 was absent in mock-challenged cells, but was readily detected 2 h after bacterial challenge in a dose-dependent manner (Fig. 5B). In the absence of STING we barely detected any phosphorylated TBK-1, whilst in the absence of MAVS a reduction was appreciated in agreement with the reporter data. We then assessed the response to PP. As shown previously, exposure to PP triggered a lower response than that to LP and control cells showed elevated luciferase activity at 24 h post challenge (Fig. 5C). However, these were also drastically reduced in STING KO cells and partially reduced in the absence of MAVS (Fig. 5C). Accordingly, levels of phosphorylated TBK1 were also diminished in the absence of STING or MAVS (Fig. 5D). Taken together, our results demonstrate that LP and PP are sensed by mechanisms that are regulated by the intracellular sensors STING and, to a lesser extent, MAVS.

**Fig. 5.**
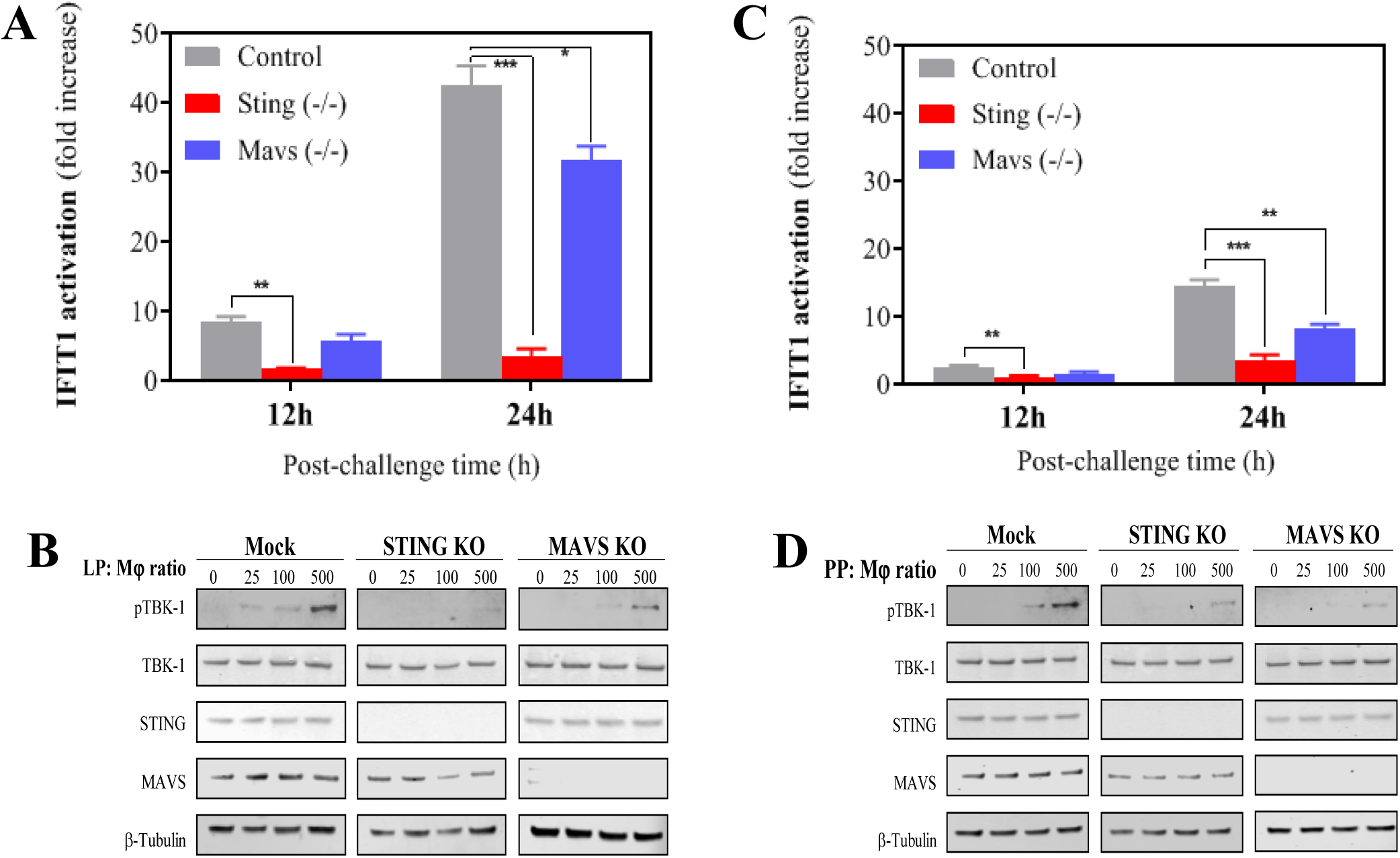
STING and MAVS sense *Lactobacillus plantarum* (LP) and *Pediococcus pentosaceus* (PP), resulting in the activation of the IFN-I associated kinase K-1. A. IFIT1 response in LP-exposed macrophages that were obtained from the pIFIT1-GLuc THP-1 cells (control, grey) and their corresponding knockouts (KO) for STING (red) and MAVS (blue). IFIT1 activation is presented as a fold increase over a non-stimulated condition after 12 and 24h of incubation using a ratio of 1 macrophage per 25 viable bacteria. For each time point, a comparative statistical analysis was carried out between the control and the knockouts using the Student t-test (*p<0.05, **p<0.01, ***p<0.005). B. Immunoblotting using whole-cell lysates from LP-challenged THP-1 macrophages against phosphorylated TBK-1 (pTBK-1), total TBK-1 (TBK-1), STING, MAVS and the housekeeping b-tubulin. The challenge was carried out for 1h using the 3 pIFIT1-GLuc THP-1 cell lines [control, STING KO and MAVS KO] and increasing ratios of macrophage per bacteria (0, 25, 100 and 500) C. IFIT1 response in PP-exposed macrophages that were obtained from the pIFIT1-GLuc THP-1 cells (control, grey) and their corresponding knockouts (KO) for STING (red) and MAVS (blue). IFIT1 activation is presented as a fold increase over a non-stimulated condition after 12 and 24h of incubation using a ratio of 1 macrophage per 25 viable bacteria. For each time point, a comparative statistical analysis was carried out between the control and the knockouts using the Student t-test (*p<0.05, **p<0.01, ***p<0.005). D. Immunoblotting using whole-cell lysates from PP-challenged THP-1 macrophages against phosphorylated TBK-1 (pTBK-1), total TBK-1 (TBK-1), STING, MAVS and the housekeeping b-tubulin. The challenge was carried out for 1h using the 3 pIFIT1-GLuc THP-1 cell lines [control, STING KO and MAVS KO] and increasing ratios of macrophage per bacteria (0, 25, 100 and 500).

### STING and MAVS contribute to IFN-I activation by LP and PP

To evaluate the influence of STING and MAVS on the ability of LP and PP to activate IFN-I responses, we determined the induction of IFN-I and IFN-I-associated genes from STING and MAVS KO macrophages exposed to LP and PP at a ratio of 25 bacteria per macrophage (Fig. 6). We first measured the relative expression of *IFN-β, MxA* and *OAS1* in the challenged KO macrophages (Fig. 6A-B). After 8 h of bacterial challenge with LP we detected a significant increase in the mRNA expression of *IFN-β* in the control cells (Fig. 6A). In both STING and MAVS KO cells, this increase was significantly lower, in particular in the STING KO cells. Similarly, the absence of STING and MAVS in cells previously exposed to LP resulted in a significant decrease in the expression levels of *MxA* and *OAS1*, two well-known ISGs. The mRNA expression of these two ISGs as well as IFN-β also decreased in STING KO and MAVS KO cells challenged with PP as compared with the control cells (Fig. 6B). However, only the reduction in IFN-β expression was found to be statistically significant. Finally, media from cells exposed to LP (the most potent inducer of IFN-I) were subjected to ISRE bioassay. In agreement with gene expression data, absence of STING and, to a lesser extent, MAVS significantly reduced the amount of biologically active IFN-I (Fig. 6C). We observed similar results with KO cells exposed to LP but the reduction in functional IFN-I was less clear and equally significant from both KO cells (Fig. 6D).

**Fig. 6.**
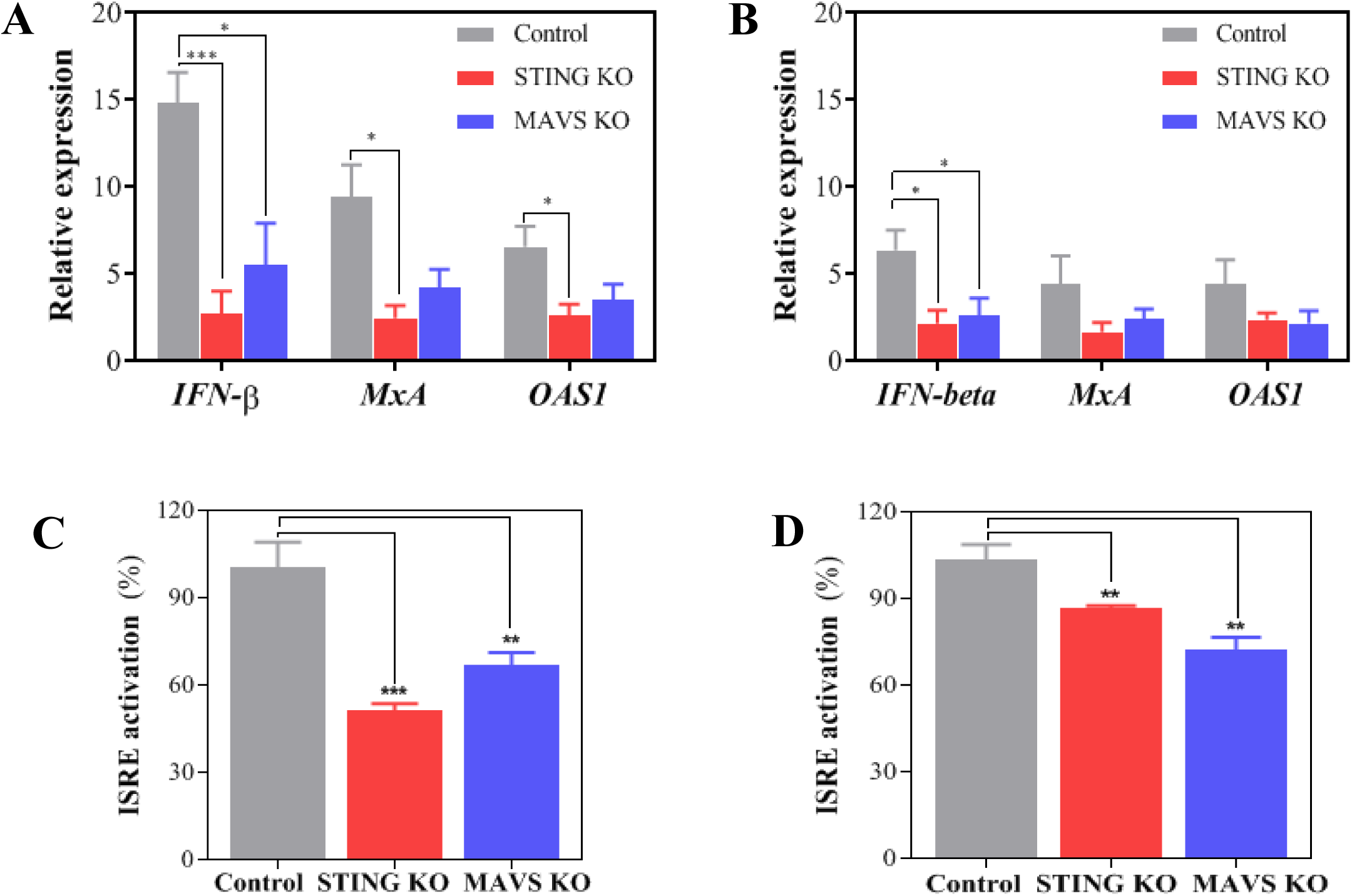
The IFN-I response of macrophages to *Lactobacillus plantarum* (LP) and *Pediococcus pentosaceus* (PP) is dependent on STING and MAVS. A. Level of relative mRNA expression of *IFN-b, MxA* and *OAS1* in pIFIT1-GLuc THP-1 macrophages (control, grey) and their corresponding knockouts (KO) for STING (red) and MAVS (blue) after exposure to LP for 8h at a ratio of 1 macrophage per 25 viable bacteria. The relative expression of each gene is presented as a fold induction following normalization with the internal control gene TBP and non stimulated THP-1 macrophages. A comparative statistical analysis was carried out between the control and the knockouts using the Student t-test (*p<0.05, ***p<0.005). B. Level of relative mRNA expression of *IFN-b, MxA* and *OAS1* in pIFIT1-GLuc THP-1 macrophages (control, grey) and their corresponding knockouts (KO) for STING (red) and MAVS (blue) after exposure to PP for 8h at a ratio of 1 macrophage per 25 viable bacteria. The relative expression of each gene is presented as a fold induction following normalization with the internal control gene TBP and non stimulated THP-1 macrophages. A comparative statistical analysis was carried out between the control and the knockouts using the Student t-test (*p<0.05, ***p<0.005). C. ISRE activation from supernatants obtained from pIFIT1-GLuc THP-1 macrophages (control, grey) and their corresponding knockouts (KO) for STING (red) and MAVS (blue) after exposure to LP for 24h at a ratio of 1 macrophage per 25 viable bacteria. The ISRE activation for each cell line was calculated as a fold increase using a pISRE-FLuc/RLuc reporter cell line after 10h of supernatant exposure and then indicated as a relative percentage considering a 100% of response for the control. A comparative statistical analysis was carried out between the control and the knockouts using the Student t-test (**p<0.01, ***p<0.005). D. ISRE activation from supernatants obtained from pIFIT1-GLuc THP-1 macrophages (control, grey) and their corresponding knockouts (KO) for STING (red) and MAVS (blue) after exposure to PP for 24h at a ratio of 1 macrophage per 25 viable bacteria. The ISRE activation for each cell line was calculated as a fold increase using a pISRE-FLuc/RLuc reporter cell line after 10h of supernatant exposure and then indicated as a relative percentage considering a 100% of response for the control. A comparative statistical analysis was carried out between the control and the knockouts using the Student t-test (**p<0.01, ***p<0.005).

## Discussion

This is the first publication reporting how human macrophages produce IFN-I in response to LAB via STING and MAVS. It is very well established that pathogenic bacteria are sensed by these cytosolic adapters to stimulate the production of IFN-I in macrophages (37). However, the role that STING and MAVS play in the recognition of beneficial bacteria such as LAB has been underexplored up to now. Our findings were generated from an initial unbiased screen using different representative LAB species. This screen showed that LAB activate IFN-I production in human macrophages in a species-dependent manner as only LP and PP were able to induce a significant induction of luciferase in THP-1-IFIT1-GLuc macrophages. This species-dependent IFTI1 activation has also been observed with other strains of LP and PP that we have recently isolated from animals (38, 39). The central dogma is that LAB activate NF-κB via TLR2 (40), TLR9 (41) and Nod-like receptors (42). Our macrophage challenge with LAB species such as *Enterococcus faecalis, Lactobacillus casei, L. sakei* and *Pediococcus acidilactici* resulted in a very high NF-κB activation, as widely reported in previous publications (38, 43-45). Unlike most of the selected LAB though, neither LP nor PP significantly activated the NF-κB pathway, but both were able to induce the exogenous production of IFN-I in human macrophage-like cells and human primary phagocytes isolated from PBMCs. This is a remarkable finding considering that only a few studies have reported that LAB are capable of inducing the production of IFN-I in innate immune cells (7, 8, 46, 47).

Here, we have observed that LP and PP up-regulate the expression of CD40 in monocytes from PBMCs, whilst displaying an antagonistic effect on CD64. Other studies with PBMCs have reported that the production of IFN-β is associated with the up-regulation of CD40 (48, 49); on the contrary, the presence of this cytokine leads to the down-regulation of CD64 (36). Therefore, our results with CD40 and CD64 suggest that the interaction of LP and PP with human phagocytes induces the production of IFN-β and this IFN-β has biological impact on human immune cells. In this respect, Weiss *et al*. (35) found that LAB that induce IFN-β activation are also able to stimulate CD40, a positive convergence that we have confirmed using macrophages derived from monocytes of PBMCs. These macrophages secrete IFN-β following the phagocytosis (or binding) of LP and PP, with a significant increase at 8h post-challenge. This production peak is in agreement with the previous work by Weiss *et al*., in which other LP isolates were used to monitor IFN-β changes in stimulated murine bone-marrow derived dendritic cells (35). These different experimental conditions could indeed explain why the levels of IFN-β production that they detected (> 500 pg/mL) are much higher than those observed by us (25-50 pg/mL).

The IFN-I activation that we have recorded with LP and PP requires interaction with paghocytes, as previously reported with dentritic cells stimulated with other LAB species (7). The viability of LP and PP was crucial to stimulate the IFN-I production in macrophages, and also that both LP and PP bind and/or are phagocytosed by human monocytes and macrophages. Alive cells of LP and PP trigger an endogenous IFN-I production that is significantly higher than that observed with inactivated cells. Based on these findings, we believe that the phagocytic intake of viable cells of LP and PP is essential to observe a good IFN-I response, which suggests a predominant role of cytosolic sensors such as STING and MAVS on the recognition of both bacterial species. In general, macrophages exposed to heat-killed (or inactivated) bacterial cells activate TRL/NOD-like receptors-dependent pathways, whereas live bacteria activate other pathways that require phagocytosis, proteolytic bacterial degradation and phagolysosomal membrane destruction, leading to the release of bacterial nucleic acids into the cytosol (50).

The cytosolic adapters STING and MAVS sense the presence of bacterial DNA or RNA in the cytosol, resulting in the production of IFN-I via a pathway dependent on the phosphorylation of TBK1 (21, 22). Our TBK-1 immunoblotting assays and the transcriptional analysis on the mRNA expression of *IFN-β* have showed that the intracellular presence of PP and LP in THP-1 macrophages activate TBK-1 and the subsequent overexpression of IFN-β. Moreover, this IFN-I activation is dependent on the presence of STING and MAVS as the THP-1 cells KO for either STING or MAVS were less capable of activating TBK-1 and IFN-β. This observation was more evident with STING KO cells exposed to LP. In this respect, some studies have reported that the bacterial recognition by STING and MAVS may be dependent on the species (37); while others have demonstrated that STING is the central player in the crosstalk between DNA and RNA sensing (51). In addition, we have proved that the production of IFN-β transcripts in macrophages challenged with LP and PP leads to the secretion of biologically active IFN-β. Through the Janus kinase signal transducer and activator of transcription (JAK-STAT) pathway, extracellular IFN-β activates the IFN-stimulated gene factor 3 (ISGF3), which binds to ISREs within ISG promoters (52). Our ISRE-based biossay responded to supernatants obtained from the challenged macrophages in a STING/MAVS dependent manner, although the influence of STING was more evident as previously observed with regards to the endogenous IFN-I activation. Furthermore, the ISGs *MxA* and *OAS1* overexpressed in macrophages exposed to LP and PP, especially to LP, and this overexpression decreased in the absence of either STING or MAVS. Therefore, our experimental data reinforces the importance of gut commensals as elicitors of protective IFN-I responses via cytosolic nucleic acid-receptors, as previously reported in colitis mouse models (25, 26). Furthermore, similarly to our findings, Moretti *et al*. (27) observed that, regardless of the expression of virulence factors, live and not dead gram-positive bacteria heightened the production of IFN-I via STING in bone marrow-derived macrophages, highlighting the pivotal role that commensal bacteria and STING play in promoting immunity and survival after infection.

In this study we have demonstrated that STING and MAVS induce IFN-I production by sensing the presence of cytosolic LAB. Nevertheless, we cannot rule out other IFN-I activation routes such as TLR2/3 recognition via endosomes, as previously reported with other LAB (7, 8). Another important aspect that is worth emphasizing is the fact that the activation of IFN-I through STING and MAVS occurs in a species-dependent manner. Why macrophages are more responsive to certain LAB species such as LP and PP and how nucleic acids and/or CDNs of these species are recognized by STING and MAVS are very important questions that remain to be elucidated. The synthesis of CDNs has been described in LAB, although very little is known about their role in bacterial physiology and innate immune responses (53). In consequence, it is early to speculate whether DNA or CDNs of LAB are more or less important for STING activation. However, the evidence that LP and PP hardly activate NF-κB suggest a major influence of DNA. RECON, a cytosolic sensor that has been discovered very recently (54), antagonize STING activation by binding bacterial CDNs, resulting in NF-κB activation.

In summary, our findings highlight the underrated value of beneficial bacteria and their molecules as potential mediators of host protective INF-I responses. STING and MAVS sense DNA and RNA of commensal bacteria to mantain gut homeostasis and prevent autoimmune conditions such as inflammatory bowel disease (25, 26). The protective role of IFN-I in the gut has been revealed recently (55) and interventions with beneficial bacteria, in particular with LP, prevent respiratory tract infections (56), which is probably due to IFN-I-based antiviral responses. A better understanding of the IFN-I regulation by LP and other LAB that have a very low impact on NF-κB-mediated pro-inflammatory cytokines could be helpful in the design of future probiotic therapies through enhancing autoimmune tolerance as well as antitumoral and antiviral immunity.

## Acknowledgements

We thank Veit Hornung (University of Munich, Germany) and Greg Towers (University College London, UK) for providing reagents. We would also like to thank Sophie Brooks and Tiago Marques Pedro for their technical assistance.

## Statement of Ethics

Ethical approval was not required

## Author Contributions

JGM planned and performed experiments, carried out data analysis and prepared and edited the manuscript. BI performed experiments, aided in data analysis, and aided in preparing and editing the manuscript. TC conducted technical experiments and edited the manuscript. FME designed experiments and aided in preparing and editing the manuscript. CMdM designed and performed experiments, aided in data analysis, and prepared and edited the manuscript. All authors read and approved the final manuscript.

